# Altered germline cyst and oocyte differentiation in *Tex14* mutant mice reveal a new mechanism underlying female reproductive life-span

**DOI:** 10.1101/814384

**Authors:** Nafisa Nuzhat, Kanako Ikami, Haley Abbott, Heather Tanner, Allan C. Spradling, Lei Lei

## Abstract

In adult mammalian females, primordial follicles that form in the fetal/neonatal ovary are the only source to sustain adult ovarian function. Our previous studies revealed that during oocyte differentiation and primordial follicle formation in mouse fetal ovaries, primary oocytes form via gaining cytoplasm and organelles from sister germ cells that are connected to them by intercellular bridges within germline cysts. To better understand the role of intercellular bridges in oocyte differentiation, we analyzed mutant females lacking *testis-expressed 14 (Tex14)*, a gene involved in cytokinesis and bridge formation. In *Tex14*^*-/-*^ fetal ovaries, germ cells divide to form a reduced number of cysts in which sister germ cells are still connected via syncytia or fragmented cell membranes, rather than normal intercellular bridges. Compared with wildtype cysts, *Tex14*^*-/-*^ cysts fragment at a higher frequency and produce a greatly reduced number of primary oocytes with highly precocious cytoplasmic enrichment and enlarged volume. By contrast, *Tex14*^*+/-*^ germline cysts are less fragmented and generate primary oocytes that are smaller than wild type. Interestingly, enlarged *Tex14*^*-/-*^ primary oocytes are much more stable than wild type oocytes and more efficiently sustain folliculogenesis, whereas undersized *Tex14*^*+/-*^ primary oocytes turn over at an accelerated rate. Our observations directly link the nature of fetal germ cell connectivity to cytoplasmic enrichment during oocyte differentiation and to oocyte developmental potential in the adult ovary. Our results imply that the duration of adult ovarian function is strongly influenced by the number of primary oocytes acquiring highly enriched cytoplasm during oocyte differentiation in fetal ovaries, rather than just by the size of the primordial follicle pool.

## Introduction

In mammals, primordial follicles containing a primary oocyte enclosed by a layer of pregranulosa cells, constitute the primitive follicular stage in the adult ovary. Most primordial follicles in the adult ovary remain quiescent, while a small fraction is periodically recruited to undergo further development to sustain folliculogenesis and egg production. A longstanding mystery of mammalian oogenesis is the extremely low efficiency in maintaining and utilizing primordial follicles. For example, in humans, a neonatal ovary contains ∼1 million primordial follicles. However, an average female only ovulates ∼500 mature eggs throughout her reproductive life-span. Moreover, primordial follicle number continuously declines in the adult ovary due to follicle recruitment and atresia. A rapid loss of primordial follicles after age 40 results in menopause in women, when less than 1000 quiescent primordial follicles persist in the ovary and folliculogenesis halts permanently (1) (2) (3) (4). Despite the central role played by primordial follicles in ovarian function, why only a tiny proportion of primordial follicles successfully complete folliculogenesis remains unknown.

Our previous studies on the mechanism of primary oocyte differentiation suggest that newly formed primordial follicles already differ in developmental potential. That is because the multiple fetal germ cells that contribute to each primary oocyte vary in number and behavior. The process of primary oocyte formation begins when newly arrived PGCs proliferate with incomplete cytokinesis from embryonic day 10.5 (E10.5) to E14.5. Germ cells derived from the same PGC remain connected by intercellular bridges, forming germline cysts. As germ cells divide, germline cysts fragment into smaller cysts of various sizes. On average, each PGC produces five cyst fragments, each containing six germ cells in E14.5 ovaries. The bridges enable germ cells to exchange signaling molecules, so that germ cells within a cyst undergo synchronized mitosis and meiosis (5) (6). Moreover, intercellular bridges enable cytoplasmic exchange between sister germ cells. By electron microscopy (EM), mitochondrial and endoplasmic reticulum (ER) were observed in the lumen of intercellular bridges joining two E14.5 germ cells (7). As oocyte differentiation progresses, intercellular bridges detach from the cell membrane, which enlarges the openings between sister germ cells around E17.5. The larger connections probably facilitate transport of large organelles, such as Golgi complexes. Cytoplasmic transport within cysts enable ∼20% of the E14.5 germ cells to differentiate into primary oocytes containing an enlarged cytoplasmic volume and organelle content by postnatal day 4 (P4), while the remaining fetal germ cells undergo apoptosis after donating these materials (8). From E14.5 to P4, germ cells that become primary oocytes increase their cell volume four-fold on average. Interestingly, individual primary oocytes in the P4 ovary still vary in cell volume, suggesting that primary oocytes acquire different amounts of cytoplasm and organelles within cysts during differentiation (8). This observation is consistent with the fact that germline cysts vary in cell number by the time germ cell division ceases at E14.5. Mouse mutants with altered germline cyst fragmentation patterns represent a potentially powerful tool for testing the relationship between cyst size, primary oocyte volume and an oocyte’s potential to undergo folliculogenesis and generate a viable egg.

During mammalian gametogenesis, intercellular bridges are found in fetal ovaries, fetal testes, and adult testes (9) (10) (11) (12) (13). Studies in adult mouse testes demonstrate that the Testis-expressed 14 (TEX14) protein is essential for blocking cytokinesis and converting transient midbody rings into stable intercellular bridges (14) (15) (16). *Tex14* mutant testes are smaller in size but contain normal spermatogonia stem cells. Stable intercellular bridges are absent in postnatal testes, where the first wave of spermatogenesis halts before the completion of the first meiotic division (13). *Tex14* mutant females are fertile, but their ovaries contain fewer oocytes compared to wildtype females. Stable intercellular bridges are not observed in *Tex14* mutant neonatal ovaries, although germ cells still form clusters (17). Due to technical difficulties, important questions, such as whether ovarian fetal germ cells still form germline cysts and remain connected without stable intercellular bridges, and whether cytoplasmic transport during oocyte differentiation continues to takes place in *Tex14* mutant fetal ovaries remain to be addressed.

In the present study, by using single-PGC lineage tracing and EM, we report that *Tex14*^*-/-*^ female germ cells form syncytial cysts or cysts with fragmented cell membrane without normal intercellular bridges, and produce a greatly reduced number of primary oocytes of increased size. Reduced amounts of TEX14 protein in *Tex14*^*+/-*^ germ cells lead to reduced cyst fragmentation and an increased number of primary oocytes that are smaller than normal. These undersized *Tex14*^*+/-*^ primary oocytes turn over at an accelerated rate. In contrast, the enlarged *Tex14*^*-/-*^ primary oocytes are much more stable than wild type, and more efficiently sustain folliculogenesis, partially compensating for their reduced number. In summary, our results reveal a new mechanism underlying female reproductive life span. Adult ovarian function is not simply determined by the size of primordial follicle pool, but is strongly influenced by the number of primary oocytes whose cytoplasm has become highly enriched by intercellular transport during oocyte differentiation in fetal ovaries.

## Results

### Formation of germline cysts lacking stable intercellular bridges in *Tex14*^*-/-*^ gonads

To directly uncover whether or not PGCs divide to form germline cysts in the absence of TEX14 protein, we first conducted single-cell lineage tracing in mutant mice. Because only one or two widely separated PGCs are labeled in each fetal gonad, this approach allows the progeny cells generated from individual PGCs to be followed despite cyst fragmentation and association with cyst fragments from separate PGCs (18) (6). After a single low dose of tamoxifen injection at E10.5, fetal ovaries and testes were collected at E14.5 when mitotic germ cell division has been completed, and stained with a GFP antibody to identify the lineage-labeled germ cells in the gonad. In *Tex14*^*+/+*^ ovaries, clustered YFP^+^ cyst germ cells were readily observed (Fig 1A). *Tex14*^*+/+*^ E14.5 cyst germ cells each have intact cell membrane (stained by an antibody to Na^+^K^+^-ATPase) and are connected by TEX14-positive intercellular bridges (Fig 1B). By EM, these stabilized intercellular bridges can be morphologically recognized joining adjacent germ cells (Fig 1C,C’). Lineage-labeled clustered cyst germ cells were found in the *Tex14*^*-/-*^ ovaries at a similar frequency as in wildtype gonads (Fig 1D, S1 Fig). The connectivity of *Tex14*^*-/-*^ lineage-labeled germ cells were shown by the lack of cell membrane (labeled by the antibody to Na^+^K^+^-ATPase) between sister germ cells (Fig 1E), and synchronized mitosis and meiosis (Fig 1F,G). To further characterize the germ cell connectivity between *Tex14*^*-/-*^ cyst germ cells at the ultracellular level, we examined germ cell morphology in E14.5 *Tex14*^*-/-*^ ovaries by EM (Fig 1H-K). E14.5 *Tex14*^*-/-*^ germ cell clusters without somatic cells in between were observed frequently. Typical intercellular bridges that were observed in *Tex14*^*+/+*^ ovaries (Fig 1C) were not found in E14.5 *Tex14*^*-/-*^ ovaries. Clustered germ cells (n=130) were found to be connected by three means: 1) 36.2% of the germ cells were connected due to fragmented membrane. Organelles were observed between the fragmented membranes (Fig 1H-I). 2) 10.8% of the germ cells formed rosette syncytia, where germ cell membranes merge at the center of the syncytia. Centrosomes of the germ cells often located at the center of the syncytia where membrane merges (Fig 1J). 3) 6.15% of the germ cells were connected via a small membrane gap between two adjacent germ cells with intact cell membrane, which may result from the failure of forming/stabilizing intercellular bridges (Fig 1K-K’’). Clustered germ cell clones were also found in E14.5 *Tex14*^*-/-*^ fetal testes (Fig 1L,M). Typical intercellular bridges were not observed by EM as well.

**Fig 1.**
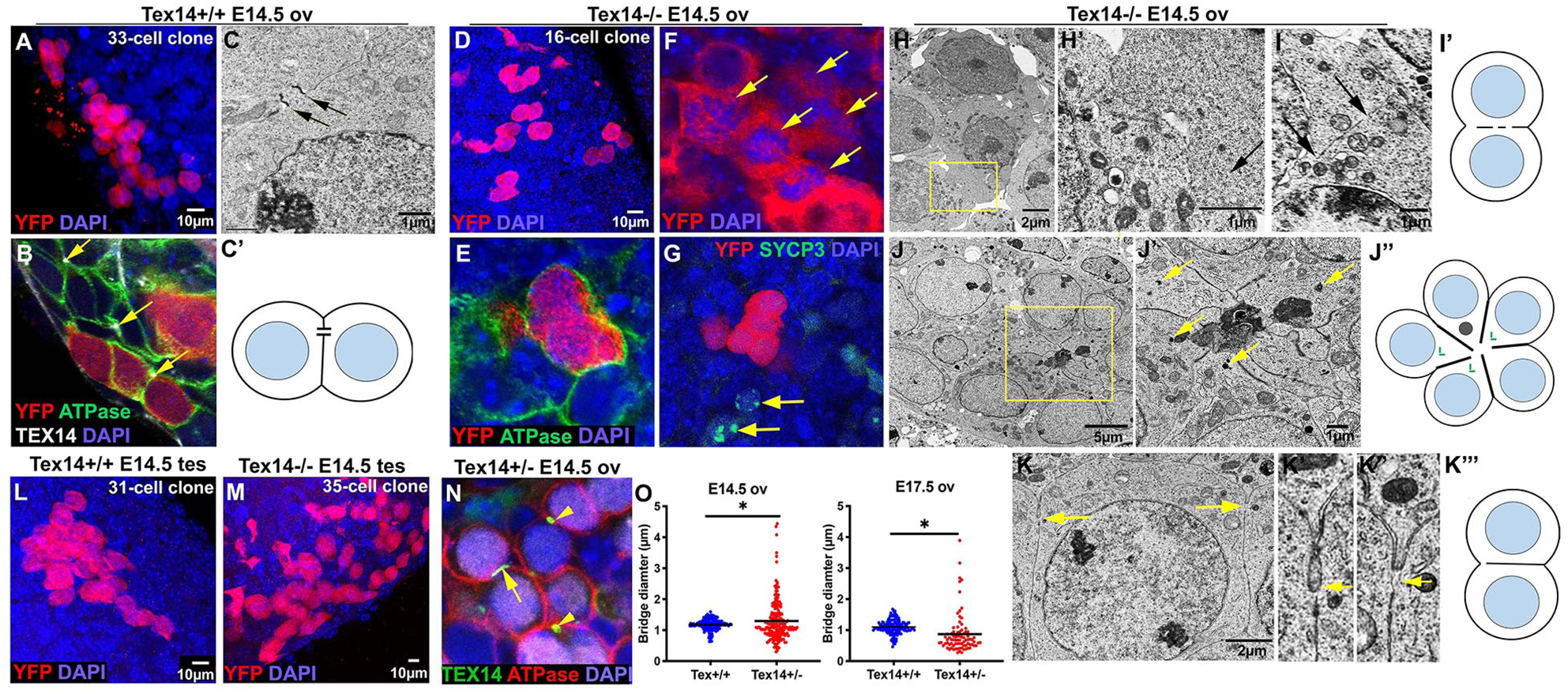
Defects in intercellular bridges in *Tex14* mutant germline cysts. (A) A YFP^+^ germ cell clone containing germline cyst fragments revealed by single-cell lineage tracing in the E14.5 Tex14^+/+^ ovary. (B) Immuno-staining showing Tex14^+/+^ cyst germ cells are connected by intercellular bridges (arrows) located at the end of the cell-cell interface. (C) An electron microscopic image and a diagram (C’) showing an intercellular bridge with cell membrane attached to both side of the bridge. (D) A YFP^+^ germ cell clone containing germline cyst fragments in the E14.5 Tex14^-/-^ ovary. (E) Immuno-staining showing Tex14^-/-^ clustered cyst germ cells with no detectable cell membrane and intercellular bridges between two cyst germ cells. (F) Synchronized mitosis revealed by clustered, dividing Tex14^-/-^ germ cells (arrows). (G) Cyst germ cells are in synchronized in entering meiosis. Among YFP^+^ cyst germ cells, none are in meiosis. Two nearby germ cells (arrows) are in meiotic division. (H-K’’’) Germ cell connectivity defects observed in E14.5 Tex14^-/-^ fetal ovaries. Clustered germ cells were found frequently in the E14.5 Tex14^-/-^ fetal ovary with fragmented cell membrane that leaves opening (arrow) between two germ cells (H-I’); (J-J’’) Tex14^-/-^ germ cells form a rosette structure where cell-cell membrane form junctions and centrosomes (arrows) are near the open area in the center of the rosette. (K-K’’’) Three Tex14^-/-^ germ cells are connected with narrow opening (arrows) at the end of the shared cell membrane. (L) A YFP^+^ germ cell clone containing a germline cyst in the E14.5 Tex14^+/+^ testis. (M) A YFP^+^ germ cell clone containing many cyst fragments in the E14.5 Tex14^-/-^ testis. (N) Fragmented TEX14^+^ intercellular bridges (arrow) and regular circular intercellular bridges (arrow heads) were observed in Tex14^+/-^ E14.5 ovaries. (O) Measurements of TEX14^+^ bridge diameters in E14.5 and E17.5 Tex14^+/-^ ovaries. F-test was used to compare the variances between *Tex14*^*+/+*^ and *Tex14*^*+/-*^ bridges.

TEX14^+^ bridges were found in *Tex14*^*+/-*^ ovaries (Fig 1N, arrowheads). However, compared with bridges in *Tex14*^*+/+*^ ovaries, fragmented bridges (Fig 1N, arrow) were observed at a higher frequency. We measured the diameter/largest cross section of the TEX14^+^ bridges in *Tex14*^*+/+*^ and *Tex14*^*+/-*^ ovaries at E14.5 and E17.5. Bridges in E14.5 *Tex14*^*+/+*^ ovaries measured 1.2 ⍰m on average, ranging from 0.6 to 1.6 ⍰m. However, bridges in E14.5 *Tex14*^*+/-*^ ovaries showed a much wider size spectrum, with an average of 1.3 ⍰m while ranging from 0.3 to 4.4 ⍰m (Fig 1O). Increased size dispersion was observed in E17.5 ovaries as well; average bridge diameter was 1.1 ⍰m in *Tex14*^*+/+*^ vs. 0.87 ⍰m in *Tex14*^*+/-*^ (Fig 1O). This observation suggests that TEX14 is haploinsufficient for stabilizing intercellular bridges (14). Reduced bridge stability may underlie the differences between *Tex14*^*+/+*^ and *Tex14*^*+/-*^ cysts. The fact that both enlarged and condensed bridges were observed in *Tex14*^*+/-*^ ovaries suggests that the tension at the midbody ring may go both expansion and contraction.

### Changes in cyst formation and fragmentation due to defective intercellular bridges

To characterize the properties of germline cysts formed with normal intercellular bridges in *Tex14*^*+/+*^ gonads, unstable intercellular bridges in *Tex14*^*+/-*^ gonads, and absent intercellular bridges in *Tex14*^*-/-*^ gonads, we analyzed the number of cyst fragments in each clone and the number of germ cells in each cyst fragment in E14.5 ovaries and testes. On average, each PGC produces clones of 30.3±19.4 cells in *Tex14*^*+/+*^, 28.4±12.0 cells in *Tex14*^*+/-*^ovaries and 14.6±6.8 cells in *Tex14*^*-/-*^ ovaries (Fig 2A). Compared with *Tex14*^*+/+*^ cysts, *Tex14*^*+/-*^ cysts fragmented less frequently, producing larger cyst fragments. *Tex14*^*-/-*^ clones contained about the same number of cyst fragments as *Tex14*^*+/+*^ clones, but on average each *Tex14*^*-/-*^ cyst fragment contained fewer germ cells (Fig 2B, C). We further profiled the distribution of the cyst fragment sizes and found that cyst fragments in *Tex14*^*+/-*^ ovaries had a higher chance of being 2-cell, 4-cell, 8-cell and 16-cell cyst fragments, i.e. cysts tended to remain connected, thus undergoing synchronized mitosis to produce cyst fragments at the size of power of 2. This observation is consistent with the result that *Tex14*^*+/-*^ cysts fragmented less frequently. Cyst fragments in *Tex14*^*-/-*^ E14.5 ovaries were smaller in size with most of them having between 2 to 8 cells. The ratio of single germ cells was slightly increased in *Tex14*^*-/-*^ E14.5 ovaries (30.9% in *Tex14*^*+/+*^ vs.37.4% in *Tex14*^*-/-*^)(Fig 2D). Germ cell clones observed in E14.5 testes of three genotypes were about the same size (Fig 2E), but these cysts fragmented very differently. *Tex14*^*+/-*^ cysts fragmented less and each contained more germ cells, similar to the fragmentation pattern observed in *Tex14*^*+/-*^ ovaries. *Tex14*^*-/-*^ cysts fragmented at a much higher rate, thus a greater number of cyst fragments at reduced size were observed compared with *Tex14*^*+/+*^ cysts (Fig 2F-G). Notably, the ratio of single cells increased significantly in *Tex14*^*-/-*^ male cysts, suggesting that male germline cysts are fragmented at a higher frequency when TEX14 protein is absent (Fig 2H).

**Fig 2.**
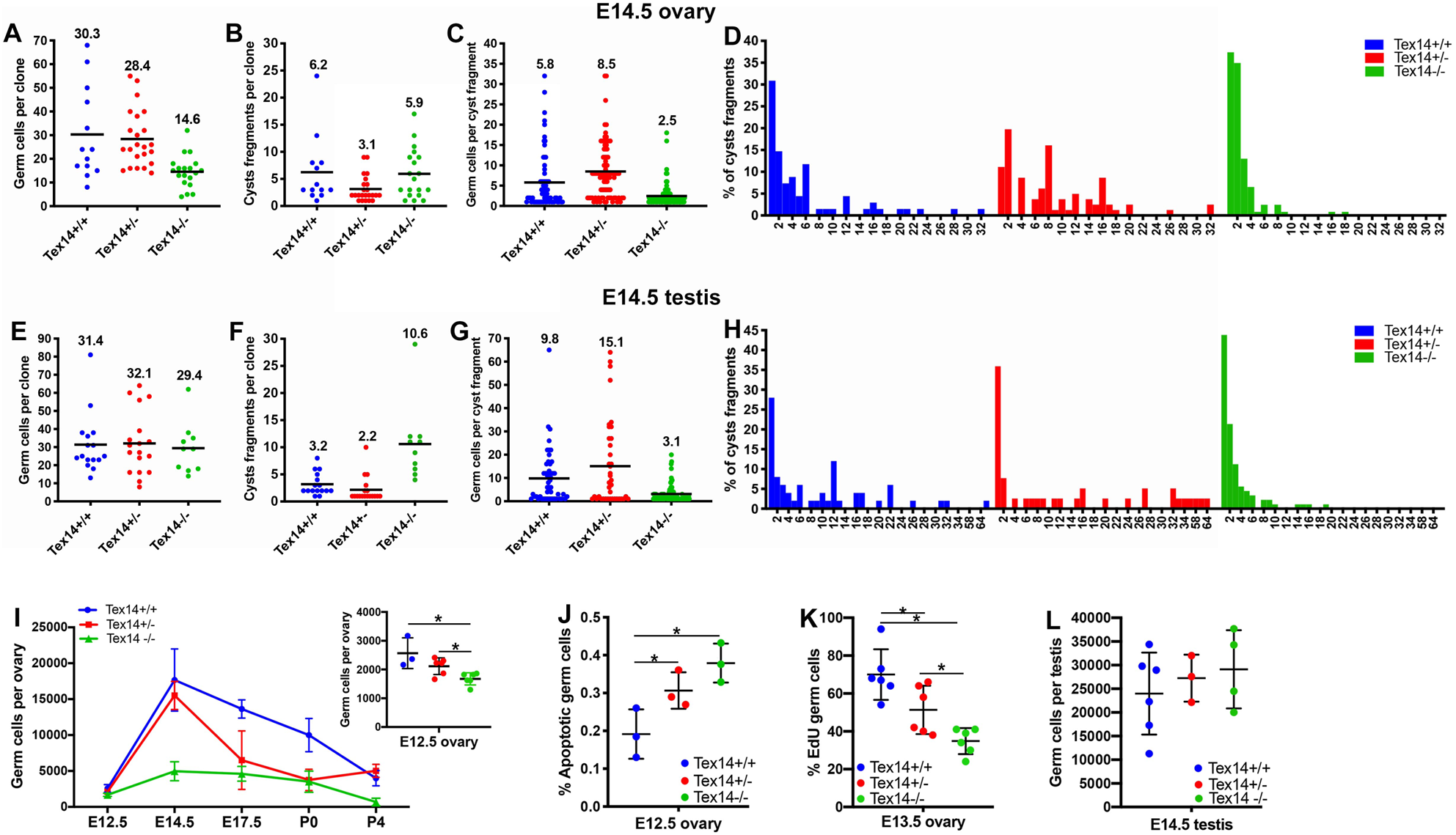
Altered germline cysts formation and fragmentation in *Tex14* mutants. Numbers of germ cells in each lineage labeled clone in E14.5 ovaries (A) and testes (E). Numbers of cyst fragments in each clone in E14.5 ovaries (B) and testes (F). Numbers of germ cells in each cyst fragment in E14.5 ovaries (C) and testes (G). The distribution of cyst fragments by size in E14.5 ovaries (D) and E14.5 testes (H). (I) Changes in germ cell number during oocyte differentiation. (J) Percentage of cleaved-Parp positive germ cells in E12.5 ovaries. (K) Percentage of EdU positive mitotic germ cells in E13.5 ovaries. (L) Number of germ cells in E14.5 testes.

To elucidate the change in germ cell numbers during oocyte differentiation from E12.5 to P4, we further quantified germ cell numbers in the fetal and neonatal ovary. In all three genotypes, fetal germ cell number per ovary reached maximum at E14.5 and began to decline afterwards. On average 24.6%, 36.5%, and 16.2% of the E14.5 fetal germ cells differentiated into primary oocytes in *Tex14*^*+/+*^, *Tex14*^*+/-*^, and *Tex14*^*-/-*^ respectively. Although the difference in germ cell number per ovary among the three genotypes started on E12.5, it increased significantly from E12.5 to E14.5. An E14.5 *Tex14*^*-/-*^ ovary only contained 28% of the germ cells as a *Tex14*^*+/+*^ ovary (Fig 2I). The decreased germ cell number in E14.5 *Tex14*^*-/-*^ ovaries is a combined result of increased germ cell apoptosis, measured as the percentage of cleaved-PARP positive cells (Fig 2J), and significantly reduced germ cell mitotic rate, measured as the percentage of EdU positive cells (Fig 2K). Interestingly, a defect in germ cell mitosis was only observed in mutant fetal ovaries, but not in testes. Germ cell numbers were comparable in E14.5 testes of the three genotypes (Fig 2L).

### Precious cytoplasmic enrichment in *Tex14*^*-/-*^ germline cysts

During oocyte differentiation in wildtype ovaries, as cytoplasmic transport within the germline cyst progresses, in the germ cell with enriched organelles, Golgi complexes gradually enucleate into a spherical structure. This spherical Golgi complex is a major component of the Balbiani body in mouse primary oocytes (8). In E14.5 *Tex14*^*+/+*^ germ cells, a linear-shaped Golgi complex was observed by EM and immunostaining of the Golgi-specific protein GM130 (Fig 3A, B). As oocyte differentiation ceases in P4 ovaries, a spherical Golgi complex was found in most *Tex14*^*+/+*^ primary oocytes (Fig 3C-D). By contrast, in E14.5 *Tex14*^*-/-*^ germ cells, where stable intercellular bridges were absent, clustered Golgi complexes were observed by using EM and immunostaining, suggesting that cytoplasmic enrichment takes place preciously in *Tex14*^*-/-*^ cysts (Fig 3 E-F). This observation was further confirmed by the measurement of germ cell diameters during oocyte differentiation. *Tex14*^*-/-*^ germ cells were larger in mean diameter starting from E12.5; in contrast, the average diameter of *Tex14*^*+/-*^ cells was smaller than *Tex14*^*+/+*^ cells. These relative differences in germ cell size remained during oocyte differentiation (Fig 3G). On average, *Tex14*^*-/-*^ primary oocytes were 1.4 times bigger than *Tex14*^*+/+*^ primary oocytes, and *Tex14*^*+/-*^ primary oocytes were only 53% of the volume of *Tex14*^*+/+*^ primary oocytes (Fig 3H).

**Fig 3.**
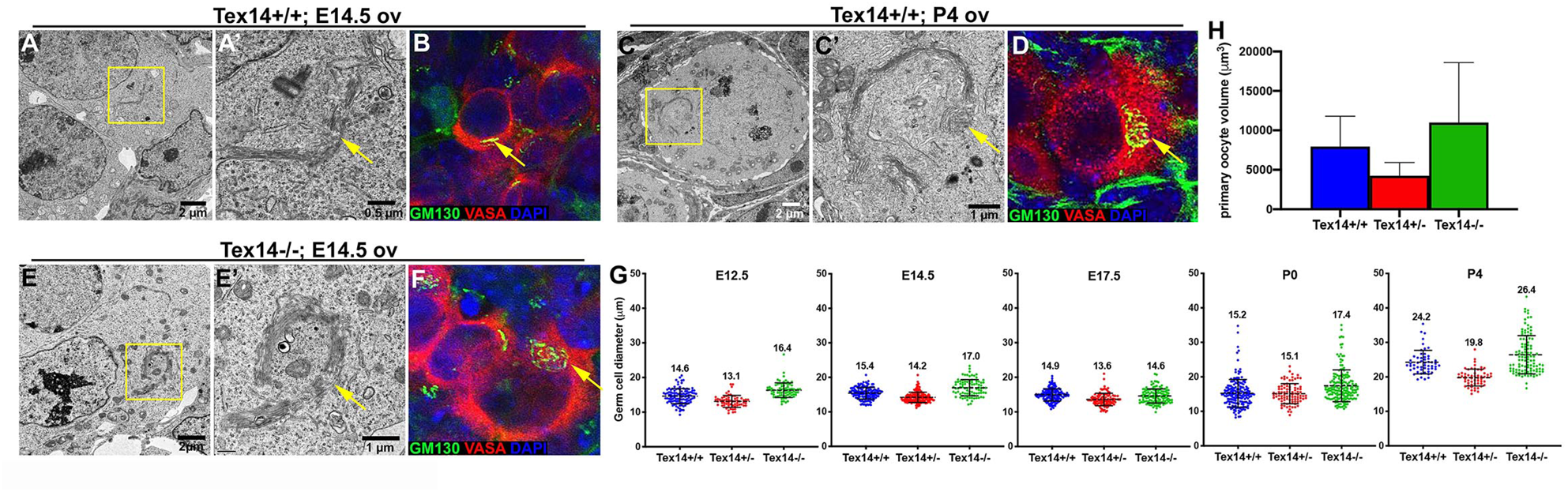
Precocious cytoplasmic enrichment during oocyte differentiation in *Tex14*^*-/-*^ ovaries. (A-B). Morphologically lineal Golgi complex was observed in the germ cells of E14.5 *Tex14*^*+/+*^ ovaries by EM (A, A’) or immuno-antibody staining (B). (C-D) In the primary oocytes of P4 *Tex14*^*+/+*^ ovaries, spherical Golgi complex was observed by EM (C, C’) and immuno-antibody staining (D). (E-F) In the germ cells of E14.5 *Tex14*^*-/-*^ ovaries, spherical Golgi complex was observed by EM (E, E’) and immuno-antibody staining (F). (G) Changes in germ cell diameter during oocyte differentiation in wildtype and *Tex14* mutant ovaries. (H) Average volumes of primary oocytes in wildtype and *Tex14* mutant P4 ovaries.

### Dynamics of folliculogenesis in adult ovaries

A previous study found that *Tex14* mutant female mice are fertile and that females of the three genotypes produced roughly the same size litters during the first six months (17). Here, we observed significant differences in the number of primordial follicles and average primary oocyte size in P4 ovaries of the three genotypes (Fig 4A). These results raised the question of whether differential utilization and/or survival of primordial follicles compensate for their numerical differences. We first quantified follicles at primordial, primary and later (any follicles beyond secondary follicle) stages by using a consistent follicle quantification approach (19) (20) (21). At P4, when primordial follicle formation is complete, a *Tex14*^*-/-*^ ovary contained only 1/6 of the primordial follicles, 1/3 of the primary follicles, and ½ of the later stage follicles of those in a *Tex14*^*+/+*^ ovary. A *Tex14*^*+/-*^ ovary contained 1.3 times more primordial follicles, 1.6 times more primary follicles, and 2.9 times more later stage follicles compared with a *Tex14*^*+/+*^ ovary (Fig 4A). However, at 1 month of age (sex maturity), *Tex14*^*-/-*^ ovaries contained ∼¼ of the primordial follicles of that in a *Tex14*^*+/+*^ovary, and comparable numbers of primary and later stage follicles as *Tex14*^*-/-*^, *Tex14*^*+/-*^, and *Tex14*^*+/+*^ ovaries. By 4 months, *Tex14*^*+/-*^ and *Tex14*^*-/-*^ ovaries contained comparable numbers of primordial, primary and later stage follicles, and significantly less than those follicles in *Tex14*^*+/+*^ ovaries (Fig 4B). Although there was no significant difference in the number of later-stage atretic follicles of three genotypes, the percentage of atretic follicles (later stage follicles containing C-PARP+ granulosa cells) was the highest in 4-month *Tex14*^*+/-*^ ovaries (Fig 4C). *Tex14*^*+/-*^ ovaries contained more corpora lutea (7.2±2.9) than those in *Tex14*^*+/+*^ (2.8±2.4) and *Tex14*^*+/-*^ (0.8±1.2) ovaries (Fig 4D). In 8-month ovaries, the size of primordial follicle pool in *Tex14*^*+/-*^ and *Tex14*^*-/-*^ were much smaller than the *Tex14*^*+/+*^ ovary (*Tex14*^*+/+*^: 939±296; *Tex14*^*+/-*^: 263±239; *Tex14*^*-/-*^: 42±19). *Tex14*^*+/-*^ and *Tex14*^*-/-*^ ovaries also contained less primary and later stage follicles. 8 months *Tex14*^*+/-*^ ovaries contained significantly more later stage atretic follicles than that in the *Tex14*^*+/+*^and *Tex14*^*-/-*^ ovaries, and comparable numbers of corpora lutea were found in the ovaries of three genotypes (Fig 4C). Although *Tex14*^*+/-*^ ovaries started with a comparable number of primordial follicles as *Tex14*^*+/+*^ ovaries, the rate of primordial follicle loss was more than 2 times faster. From 1 month to 8 months, the half-life of the primordial follicle pool was 4.8 months in *Tex14*^*+/+*^, 2.3 months in *Tex14*^*+/-*^ and 1.7 months in *Tex14*^*-/-*^ (Fig 4B). Interestingly, compared with *Tex14*^*+/-*^, *Tex14*^*-/-*^ is more efficient in utilizing primordial follicles for sustaining folliculogenesis, the half-life of the total follicles from 1 month to 8 months was 16.9 months in *Tex14*^*+/+*^, 5.0 months in *Tex14*^*+/-*^ and 8.2 months in *Tex14*^*-/-*^ (Fig 4E).

**Fig 4.**
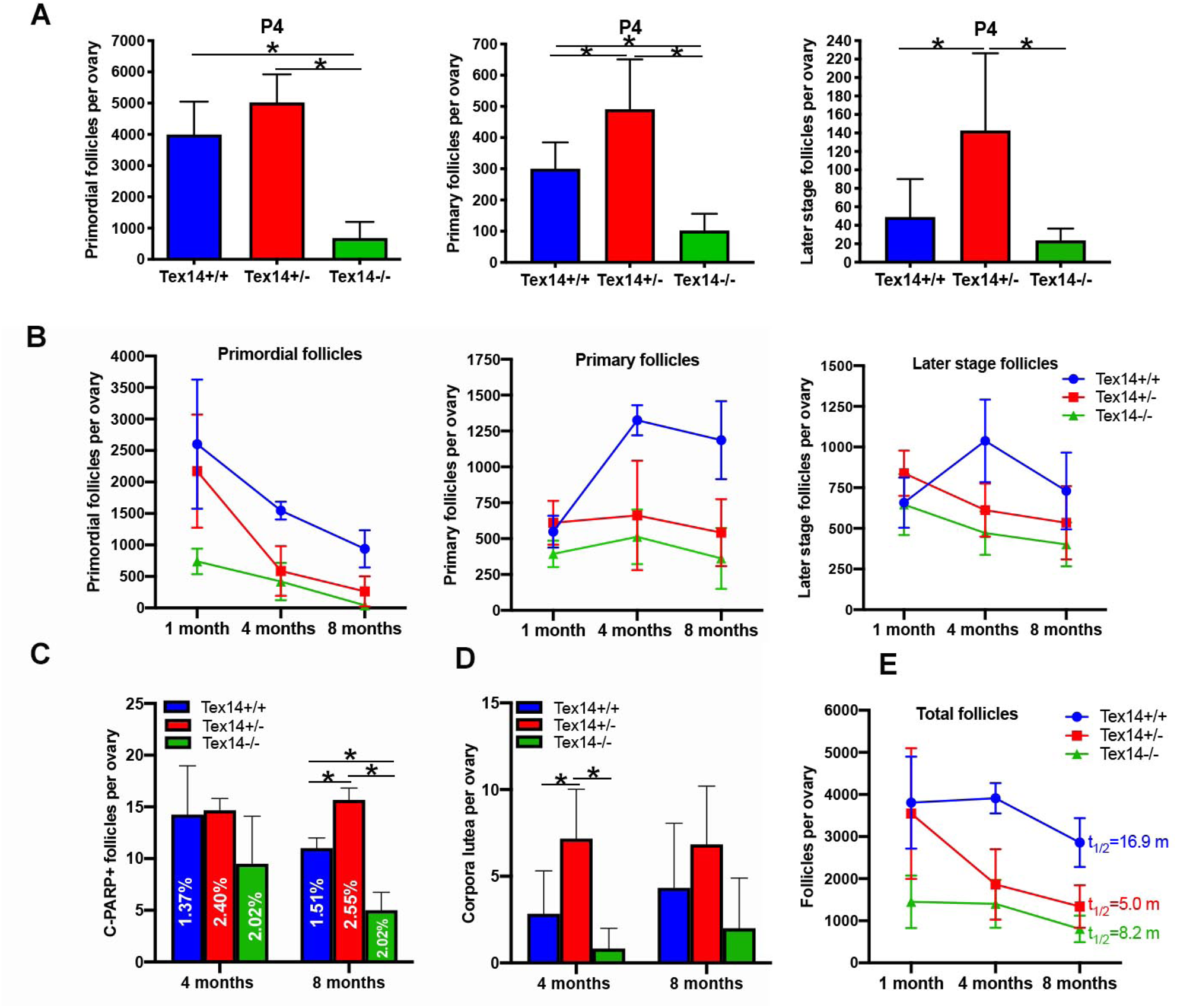
Dynamics of folliculogenesis in wildtype and *Tex14* mutant ovaries. (A) Numbers of primordial follicles, primary follicles and later stage follicles (secondary follicles and beyond) in P4 ovaries. (B) Changes in the numbers of primordial follicles, primary follicles and later stage follicles in adult ovaries from 1 month to 8 months. (C) Numbers of atretic later stage follicle in 4 months and 8 months ovaries. (D) Numbers of corpora lutea in 4 months and 8 months ovaries. (E) Changes in total follicle numbers in adult ovaries from 1 month to 8 months.

### TEX14 protein gradually accumulates on intercellular bridges during germline cyst formation

Our previous study revealed that on average, each PGC proliferate to form 2-cell or 4-cell cysts at E11.5, and 4-cell, 8-cell or 16 cell cysts by E12.5. Cyst formation gradually lose synchrony due to fragmentations and by E14.5 each PGC produces a 30-cell clone on average (6). To further characterize the mechanism underlying defective intercellular bridges and changes in germline cyst formation in *Tex14* mutants, we stained E12.5 and E14.5 ovaries with a RacGAP antibody that labeled early bridges and a TEX14 antibody that labels stable bridges (14). We found that in E12.5 ovaries, the RacGAP antibody labels significantly more bridges than the TEX14 antibody. All TEX14^+^ bridges were RacGAP^+^, however, only 51±4% of the RacGAP^+^ bridges were TEX14^+^ in E12.5 ovaries (Fig 5A-A’’). When we observed bridges from the cross section of the ring in E12.5 ovaries, RacGAP was found to distribute throughout the ring with several spots stained higher level of RacGAP protein. TEX14 was not accumulated on the ring completely yet, and appeared as several patches along the RacGAP^+^ ring (Fig 5B-B’’). This observation is consistent with the role of RacGAP in cytokinesis, and the function of TEX14 in stabilize the midbody rings into stable intercellular bridges. When we examined the pattern of RacGAP and TEX14 distribution from the lateral face of the bridges in E12.5 ovaries, RacGAP distributed on the outer rim of the bridge that associate with cell membrane, and TEX14 distributed exclusively between two RacGAP^+^ outer rims of the bridge (Fig 5C-C’’). By E14.5, the ratio of RacGAP and TEX14 double positive bridges was 92±5% in ovaries. At the cross sections of a bridge, both RacGAP and TEX14 protein distributed thought out the ring (Fig 5D-D’’). Interestingly, when observing the bridge from the lateral face, RacGAP was exclusively enriched in the inner layer of the ring, while TEX14 protein localized on the outer rim of the ring which interacts directly with the cell membrane (Fig 5E-E’’). This suggests that during the later stage of cyst formation, TEX14 replaces RacGAP and interacts with cell membrane to stabilize intercellular bridges at the certain location of the germ cell membrane in order to maintain cyst structure (Fig 5F). In E14.5 *Tex14*^*-/-*^ ovaries, RacGAP^+^ small foci located between two cells were observed occasionally, but RacGAP^+^ ring-shaped bridges were not found (S2 Fig). This observation suggests that TEX14 may not be essential for blocking cytokinesis during fetal germline cyst formation, but play a major role in stabilizing the bridges on the cell membrane. This observation of TEX14 accumulation on intercellular bridges is also consistent with timing of germ cell mitosis defect observed from E12.5 to E14.5 in *Tex14*^*-/-*^ ovaries, where cyst formation takes place prior to TEX14-mediated bridge stabilization.

**Fig 5.**
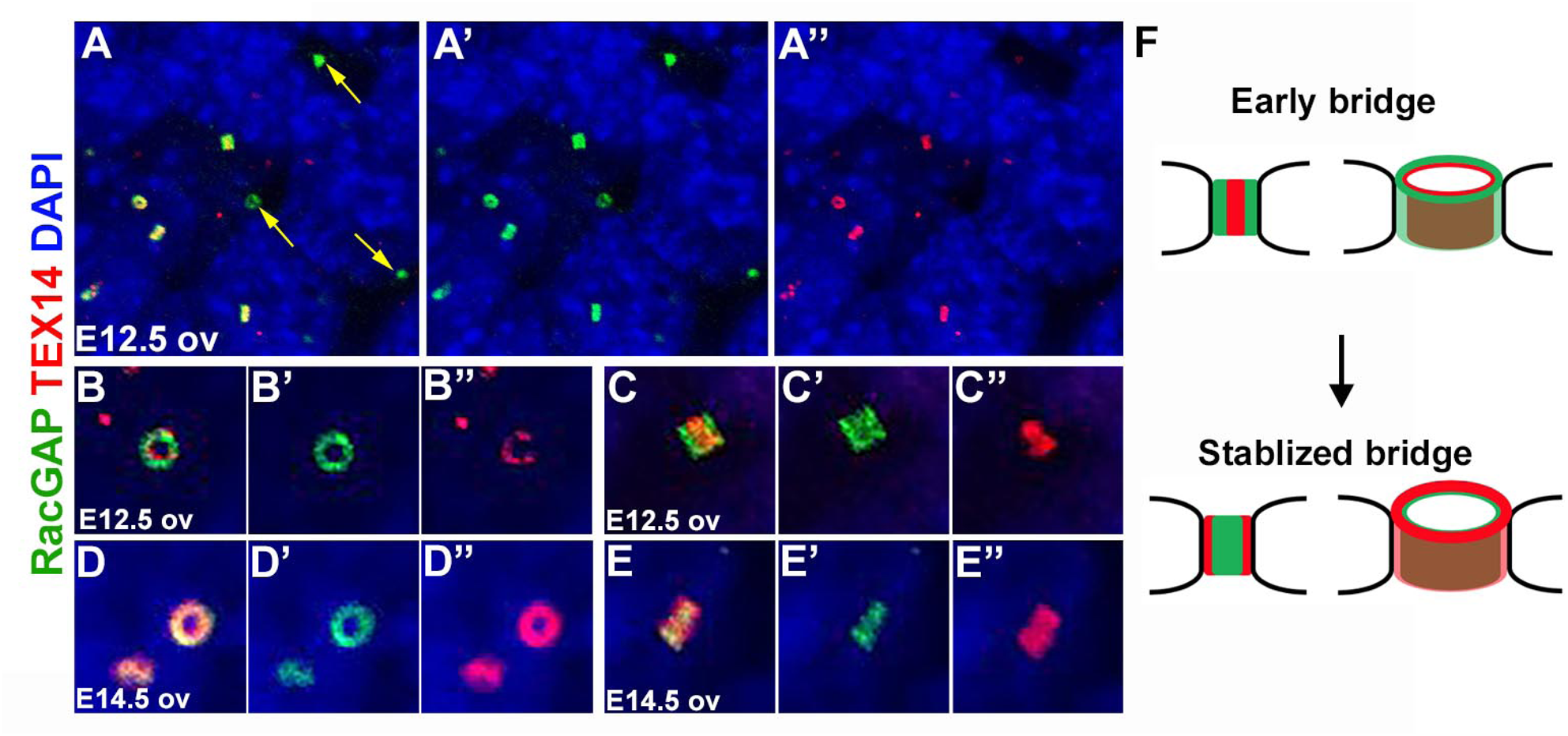
TEX14 protein gradually accumulates on RacGAP^+^ bridges during germline cyst formation. (A-A’’) In E12.5 ovaries, some RacGAP^+^ bridges are TEX14 negative (arrows). (B-B’’) A cross section of an E12.5 bridge showing TEX14 protein distributes partially on the RacGAP^+^ bridge. (C-C’’) A longitudinal section of an E12.5 bridge showing RacGAP protein locates at the outer layer of the bridge and TEX14 locates at the center of the bridge. (D-D’’) A cross section of an E14.5 bridge showing TEX14 protein completely overlap with RacGAP protein on the bridge. (E-E’’) A longitudinal section of an E14.5 bridge showing TEX14 locates throughout the bridge and RacGAP locates at the inner layer of the bridge. (F) A diagram showing the process of bridge stabilization via TEX14 accumulation on the bridge during cyst formation.

## Discussion

A lingering mystery of mammalian oogenesis is the extremely low efficiency of egg production during oogenesis. Among primordial follicles in the ovarian reserve, the fate of becoming a mature follicle vs. undergoing atresia is likely the consequence of complex interactions between intrinsic (oocyte developmental potential) and extrinsic (somatic cells and hormonal signals) factors. Our previous study demonstrated that primary oocytes in P4 ovaries vary in cell volume and Golgi complex content. This suggests that primary oocytes may collect different amount of cytoplasmic content from sister germ cells (8). The connection between cytoplasmic enrichment and primary oocyte size was also revealed by an *in vitro* experiment where cytoplasmic enrichment was blocked in cultured ovaries by treating with inhibitors to microtubule polymerization or dynein activity. Primary oocytes in these ovaries were smaller in cell size and were not able to commit to folliculogenesis in culture. This indicates that the amount of cytoplasmic content primary oocytes collect may underlie their developmental potential. How oocytes differentiate within cysts of various sizes, and whether primary oocyte developmental potential is determined by the cytoplasm they collect during differentiation are intriguing open questions. Mouse models with altered germline cyst size are ideal modes to examine the connections between cyst size, cytoplasmic enrichment and oocyte developmental potential. In the present study, the difference in cyst formation and fragmentation in the *Tex14*^*+/+*^, *Tex14*^*+/-*^, and *Tex14*^*-/-*^ mice due to the defect in intercellular bridge formation and stabilization provided us an ideal model to address these questions.

During mouse oocyte differentiation, as PGCs divide to form germline cysts from E10.5 to E14.5, cysts undergo constant fragmentation (6). Larger cyst fragments observed in *Tex14*^*+/-*^ ovaries and smaller cyst fragments in *Tex14*^*-/-*^ ovaries allowed us to tie the connection between cyst size and primary oocyte formation. In E14.5 wildtype ovaries, on average each cyst contains 6 cells, 24.6% of the E14.5 germ cells become primary oocytes by P4, and primary oocytes increase their cell volume 4.0 times from E14.5 to P4. In *Tex14*^*+/-*^ ovaries, cysts are larger in size with 8.5 cells per cyst; the efficiency of the oocyte formation is higher with 36.5%, but primary oocytes enriched less cytoplasm compared with that of wildtype. In *Tex14*^*-/-*^ ovaries, cysts fragmented to a greater extend. On average, 16.2% of the E14.5 germ cells become primary oocytes and primary oocytes enriched 6.5 folds of cytoplasm. It is compelling that large cysts produce more primary oocytes in smaller volume rather than a few primary oocytes with a greater volume, suggesting there may be a mechanism that determines the rate of oocyte production so that no primary oocytes with extra-enriched cytoplasm are produced. This result also suggests that the size of germline cysts may underlie the number of primary oocytes formed during ovarian reserve formation; cyst fragmentation observed in mouse fetal ovaries may serve as a means to produce cysts at “manageable” sizes which allows cytoplasmic enrichment to a few germ cells within a short window. Open questions, such as, whether cysts at all sizes produce primary oocytes, and how cysts fragment during oocyte differentiation will be addressed by using proper tools, such as live-imaging, to fully understand the control of primordial follicle number during ovarian reserve formation.

A surprising result we learned by analyzing follicle development in adult ovaries is that the size of primary oocytes has a strong connection with the half-life of primordial follicle pool, and that the size of primordial follicle pool does not determine the number of follicles that ultimately undergo folliculogenesis during adulthood. A *Tex14*^*-/-*^ ovary contains only 1/6 of the primordial follicles of that in a wildtype ovary. However, most of the *Tex14*^*-/-*^ primary oocytes contain greater cell volume than *Tex14*^*+/+*^ primary oocytes at P4. *Tex14*^*-/-*^ females are fertile with a comparable number of developing follicles at 4 weeks as that of the wildtype ovary. By contrast, A *Tex14*^*+/-*^ ovary contains slightly more primordial follicles at P4 initially, and majority of these primary oocytes are in smaller cell volume compared with wildtype. In adult *Tex14*^*+/-*^ ovaries, despite the larger size of the ovarian reserve, *Tex14*^*+/-*^ primordial follicles have the shortest half-life and later stage follicles undergo atresia at the highest rate. This observation suggests that the cytoplasmic content primary oocytes acquire during fetal development determines their potential to complete folliculogenesis during adulthood.

Our present study also revealed the difference in intercellular bridge formation and function between oogenesis and spermatogenesis. In adult testes, TEX14 is the key protein that stops cytokinesis and converts the midbody ring into a stable intercellular bridge (14). In the absence of TEX14, male germ cells that complete cytokinesis become individualized. Although the first cycle of spermatogenesis can initiate and progress to meiotic stages, no spermatids and only a reduced number of spermatocytes were observed in P21 *Tex14*^*-/-*^ testes due to early meiotic death (13). Our observations on the timing and localization of TEX14 protein in fetal ovaries suggests that TEX14 functions to stabilize bridges during mid to late cyst formation (E12.5-E14.5), instead of blocking cytokinesis. During early cyst formation from E10.5 to E12.5, about half of the RacGAP^+^ intercellular bridges were TEX14^-^. These TEX14^-^ bridges may enable flexibility of bridges at the cell membrane, which allows the formation of branched cysts observed in E14.5 ovaries (8). The stabilization of the bridges to the cell membrane may be achieved by changing the location of TEX14 from the inner ring to the outer ring in order to directly interacting with the cell membrane. Similar inner/outer layer of bridges and bridge stabilization have been well characterized in *Drosophila* gametogenesis. In both male and female cyst, mature ring canals consist of an outer rim closely associated with the plasma membrane and an inner rim that appears less dense in electron micrographs (22). *Drosophila* intercellular bridges also experience a maturation process during which bridge composition changes. Shortly after germ cells complete the fourth round of mitosis, phosphotyrosine epitopes accumulate at the site of the contractile ring and transform these arrested cleavage furrows into mature ring canals (23) (24). Interestingly, TEX14 protein contains a kinase domain, although kinase activity has not been observed *in vitro*. An interesting difference in intercellular bridges between *Drosophila* oogenesis and mouse oogenesis is the phenomenon of bridge expansion. During *Drosophila* oogenesis, as the large-scale cytoplasmic transport takes place, the size of the intercellular bridges expands from 1-2 ⍰m initially to ∼10 ⍰m. Mouse bridges don’t expand, but detach from the cell membrane and expand cell-cell connectivity to facilitate the large-scale cytoplasmic transport. Precocious cytoplasmic enrichment we observed in *Tex14*^*-/-*^ ovaries reveals that stable intercellular bridges also play a role in limiting the rate of cytoplasmic transport during mouse oocyte differentiation.

In summary, by characterizing the phenotypes of germ cell connectivity and germline cyst fragmentation in *Tex14* mutant mouse ovaries, we found additional evidence supporting that the transfer and accumulation of cytoplasm within cysts during primary oocyte formation is a centrally important aspect of mammalian oogenesis. Cytoplasmic transfer did not require normal intercellular bridges, but took place with the same general polarity as in wild type. Our present study uncovered the novel connections between cyst size, primary oocyte number/cell volume and the half-life of the primordial follicle pool. We demonstrated that instead of the size of ovarian reserve, the amount of the cytoplasm primary oocytes gained within cysts strongly influences their potential to commit folliculogenesis and thus the life-span of the ovarian reserve and ovarian function.

## Materials and Methods

### Mice

CAG-creER (004682) and R26R-YFP (006148) mouse strains were acquired from the Jackson Laboratory (25) (26). Tex14 mutant mouse line was generously provided by Dr. Martin Matzuk’s lab at Baylor College of Medicine (27). All mice were maintained at C57BL/6 background and housed and bred according to the protocol approved by the Institutional Animal Care & Use Committee (IACUC) at the University of Michigan or the Buck Institute for Research on Aging.

### Single-cell lineage labelling

Single-cell lineage tracing was carried out by using the protocol published in our previous study (6) (8). Briefly, a single dose (0.4 mg per 40g body weight) of tamoxifen was injected to the females (*R26R*^*YFP/YFP*^;*Tex14*^*+/-*^) mice that were plugged by male (*CAG-creER*^*+/-*^; *R26R*^*YFP/YFP*^;*Tex14*^*+/-*^) mice at E10.5. Fetal ovaries and testes were collected at E14.5 and lineage-labeled clones were revealed by YFP antibody staining.

### Whole mount immunostaining and fetal germ cell quantification

Fetal ovaries and testes were dissected in cold phosphate buffer saline (PBS) and fixed in 4% paraformaldehyde (PFA) for 1 hour at 4 degree. Tissues were washed in PBST_2_ (PBS with 0.1% Tween 20 and 0.5% Triton X-100) and incubated in primary antibodies (S1 Table) at 4 degree overnight. On the next day, tissues were washed in PBST_2_ and incubated in secondary antibodies at 4 degree overnight. The tissues were washed in PBST_2_ and incubated in DAPI at room temperature for 30 mins to stain nuclei. The stained gonads were mounted on glass microscope slides using imaging spacers and analyzed using confocal microscopy. Fetal gonads stained with anti-VASA antibody were used for germ cell number quantification. Germ cells of the largest cross section were imaged using confocal microscopy and counted by using Image-J image analysis software. The germ cell diameter was obtained by averaging two diameters at the largest cross section of the cell using Image-J. The thickness of the ovary was obtained by measure the distance from the top to the bottom of the tissue. Total number of germ cells per gonad was determined by using the following calculation: gonad thickness/germ cell diameter x germ cell number on the largest cross section.

### Follicle quantification of adult ovaries

Adult ovaries were fixed in 4% PFA overnight at 4 degree. After fixation, ovaries were washed in PBS and incubated in 30% sucrose overnight before embedded in optimal cutting temperature (OCT) compound. Serial sections were cut at 10 *µ*m of the entire ovary. Sections were stained by using VASA antibody to reveal oocytes. Follicles were counted on every fifth section. The number of follicles per ovary were calculated as follicles/section x total sections. After follicle quantification, sections were stained by using hematoxylin and eosin (H&E) for corpus luteum (CL) quantification. Based on the average size of CL (400 *µ*m), CL were counted at every 40 sections and were summed for the total number of CL in the ovary (28) (29).

### Apoptotic and mitotic germ cell assay

#### Apoptosis assay

fetal ovaries were fixed and stained by using the cleaved-PARP antibody. The number of cleaved-PARP positive cells in an ovary was quantified by counting through serial confocal sections of the ovary by using Image-J. % of apoptotic cells = % (cleaved-PARP ^+^ cells/total germ cells).

#### Proliferation assay

fetal ovaries were dissected and incubated with EdU (50 ⍰M) for 30 mins in DMEM/F12 medium with 3mg/ml bovine serum albumin. The ovaries were fixed and stained with EdU labeling kit and VASA antibody. EdU^+^ and VASA^+^ cells in a 100 *µ*m cubical area of the serial confocal images at the periphery region of the ovary were quantified by using Image-J. % of mitotic germ cells = % (EdU^+^; VASA^+^ cells/ VASA^+^ cells).

### Electron Microscopy

Electron microscopic imaging was done by Mike Sepanski at the Carnegie Institution as previously described and the Microscopy and Image Analysis Laboratory (MIL) at the University of Michigan (8).

### Statistics

All data was presented as mean±SD. Non parametric t tests were run to analyze the difference between two experimental groups. Multiple experimental groups were analyzed using one-way ANOVA. P value level of at least P<0.05 was considered to be statistically significant.

## Acknowledgements

Research reported in this paper was supported by National Institute of General Medical Sciences of the National Institutes of Health under award number R01GM126028, and the Sergey Brin Family Foundation. The authors would like to thank Dr. Hao Yan and Ronald Pandoy in the Lei lab for critical reading of the manuscript.

## Supporting information captions

**S1 Fig. Single-PGC lineage labeling efficiency.** The efficiency of single PGC-lineage labeling shown by the percentage of fetal gonads containing YFP^+^ germ cell clones in wildtype and *Tex14* mutant mouse fetal gonads.

**S2 Fig. RacGAP**^**+**^ **intercellular bridges are absent in E14.5 *Tex14***^***-/-***^ **mouse ovaries.** (A) Most intercellular bridges (arrows) are RacGAP and TEX14 positive in E14.5 *Tex14*^*+/+*^ ovaries. (B) Ring-shaped RacGAP^+^ intercellular bridges are absent in E14.5 *Tex14*^*-/-*^ ovaries, RacGAP^+^ foci were observed in the fetal germ cells occasionally.

**S1 Table. Antibodies used in the manuscript.**

