## Supplementary figures and images for "Altered germline cyst and oocyte differentiation in *Tex14* mutant mice reveal a new mechanism underlying female reproductive life-span"

### Supplemental figures

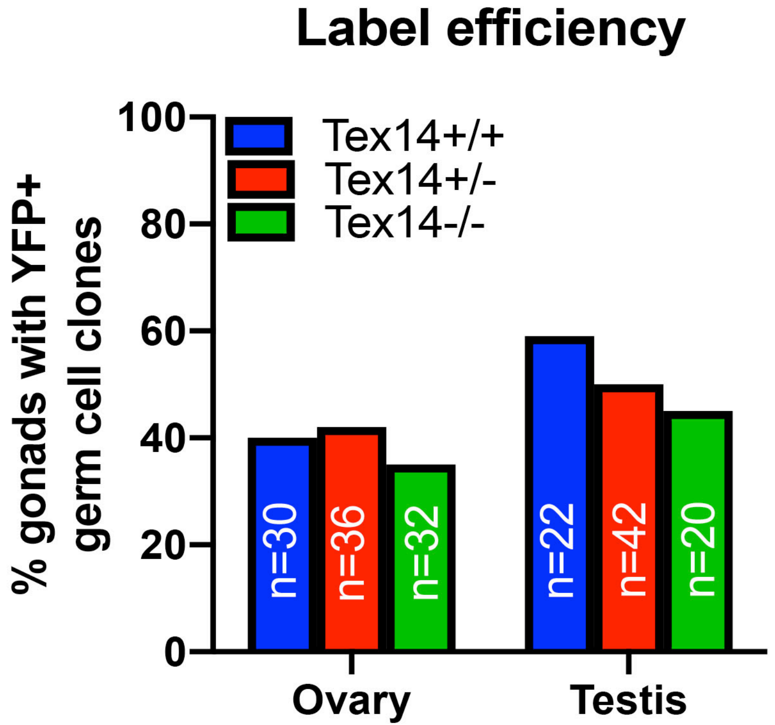

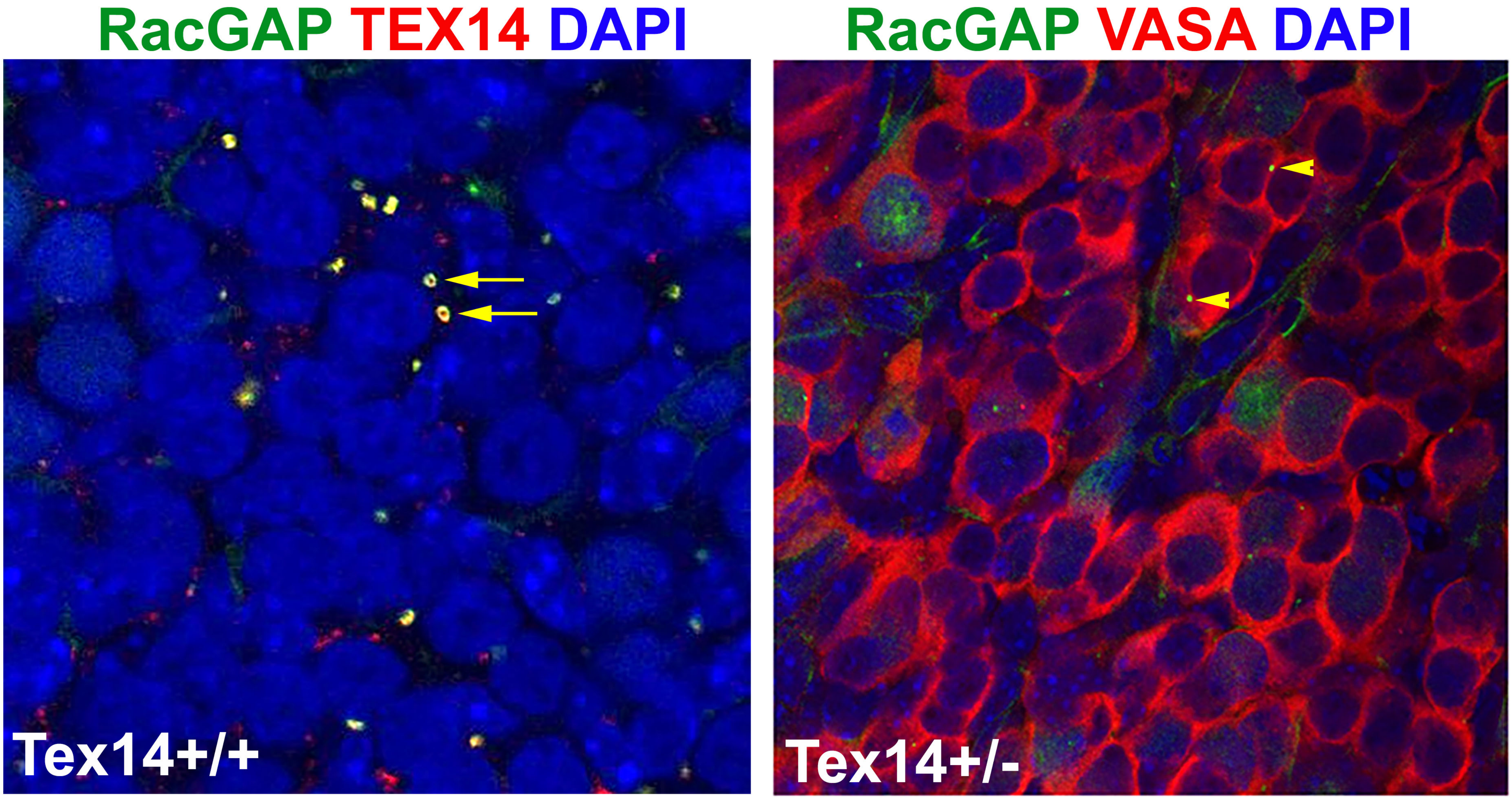
